# TDP-43 mutations-induced defects in miRNA biogenesis and cytotoxicity by differentially obstructing Dicer activity in *Drosophila* and in vitro

**DOI:** 10.1101/2024.07.10.602849

**Authors:** Xiang Long, Mengni Jiang, Yongzhen Miao, Huanhuan Du, Ting Zhang, Zhuoya Ma, Jiao Li, Chunfeng Liu, Hongrui Meng

## Abstract

Mutations in the DNA/RNA-binding protein 43 (TDP-43) can cause amyotrophic lateral sclerosis and frontotemporal dementia (ALS-FTD). As an RNA-binding protein, TDP-43 plays a diverse physiological role in RNA processing and is potentially involved in the pathological progression caused by disease mutations. However, the precise mechanisms linking RNA dysregulation and TDP-43 mutations, which propel disease progression, are not yet fully understood. Here, we demonstrate that TDP-43 and its *Drosophila* homolog, TBPH, whose mutations crucially perturb Dicer presentation, result in dysregulated miRNA profiles. Genetically modulating the expression or pharmacologically activating Dicer implies unique interactions of TDP-43 A315T and M337V with the Dicer protein, suggesting a specific mechanism of C-terminal disease mutations contributes to the pathological process. Unlike TDP-43 A315T, the M337V mutation causes the assembly of aggregates and reorganizes but functionally preserves the Dicer activity in miRNA processing. Mutations in TDP-43 disrupt miRNA biogenesis by hindering its interaction with Dicer, leading to cytotoxicity and providing mechanistic insight into the pathogenic mutations associated with ALS-FTD.

## Introduction

Amyotrophic lateral sclerosis (ALS) is a rapidly progressing disease that causes the loss of motor neurons in the brain and spinal cord, leading to skeletal muscle weakness and wasting(Brown & Al-Chalabi, 2017; Neumann *et al*, 2006). Approximately 90% of ALS cases are sporadic, while the remaining 10% are attributed to inherited mutations in genes(Chia *et al*, 2018; Goutman *et al*, 2018; Oskarsson *et al*, 2018), such as GGGGCC repeat expansions in C9orf72(Balendra & Isaacs, 2018), transactivation response element DNA/RNA-binding protein of 43 kDa (TDP-43)(Sreedharan *et al*, 2008), and fused in sarcoma (FUS)(Kwiatkowski *et al*, 2009). Many gene mutations associated with ALS are also linked to another age-related neurodegenerative disorder, frontotemporal dementia (FTD)(Ling *et al*, 2013). TDP-43 aggregation in the cytoplasm and its depletion from the nucleus of affected neurons is a notable feature in most ALS cases and approximately 45% of individuals with FTD(Igaz *et al*, 2009; Ling *et al*., 2013; Neumann *et al*., 2006; Zhang *et al*, 2009), implicating TDP-43 dysfunction in most ALS cases.

With its primary localization in the nucleus, TDP-43 modulates gene expression by participating in microRNA (miRNA) processing, transcriptional regulation, and pre-mRNA splicing(Buratti, 2015; Lagier-Tourenne *et al*, 2010). TDP-43 consists of tandem RNA-recognition motifs (RRM1 and RRM2) located near RNA splice sites and an extended C-terminal domain (CTD) rich in glycine and glutamine/asparagine(Conicella *et al*, 2016; Conlon & Manley, 2017; Lim *et al*, 2016). Most pathogenic mutations in TDP-43 are localized in the glycine-rich region and are missense mutations. These mutations result in the redistribution of the protein from the nucleus to the cytoplasm, which is commonly observed in both ALS and FTD(Rutherford *et al*, 2008). Despite extensive research on the physiological role of TDP-43 in the nucleus, understanding of its cytoplasmic RNA targets and protein interactors, as well as the impact of ALS-linked mutations on its normal functions, remains limited.

The mature miRNAs originate from endogenous hairpin-shaped transcripts in the nucleus and are then transported to the cytoplasm, where they specifically regulate gene expression by inhibition and/or degradation of target mRNA(Bartel, 2009; Schnall-Levin *et al*, 2010). Nuclear RNase III, Drosha, combines with its essential cofactor DGCR8 (also known as Pasha in *Drosophila*) to form a microprocessor complex and initiates maturation after transcription by cutting the stem-loop to release pre-miRNA(Denli *et al*, 2004; Han *et al*, 2004; Lee *et al*, 2003). After being transported to the cytoplasm, the pre-miRNA undergoes cleavage by Dicer to yield a mature miRNA of approximately 22 nucleotides. In *D. melanogaster*, DCR-1 is indispensable for miRNA biogenesis, whereas DCR-2 is responsible for generating endogenous and exogenous siRNAs, despite both enzymes possessing RNase III enzyme activity(Lee *et al*, 2004). DCR-1 interacts with Loquacious (Loqs), composed of three double-strand RNA-binding domains that regulate the efficiency of mature miRNA processing(Fukunaga *et al*, 2012; Jouravleva *et al*, 2022; Liu *et al*, 2007). Dic-2 in complex with its partner, R2D2, forms an RNA-induced silencing complex (RISC) loading complex, binds to a perfectly complementary siRNA and is sorted into AGO2(Liu *et al*, 2009; Pham *et al*, 2004). Intriguingly, miRNA can regulate multiple biological pathways by targeting one or several key genes simultaneously in mammals or fruit flies. Any disruption to this pathway could potentially have detrimental consequences on gene expression networks and cellular homeostasis.

Given the fact that dysregulation of miRNA is crucial for the maintenance of neurons by regulating genes and pathways associated with ALS-FTD(Eacker *et al*, 2009; Liu *et al*, 2012), abnormal miRNA processing is expected to be linked to the disruption of cytoplasmic activation of pertinent proteins. TDP-43 has been confirmed as a component of the nuclear microprocessor complex(Gregory *et al*, 2004), with its involvement impacting the biogenesis of a subset of miRNAs(Di Carlo *et al*, 2013; Kawahara & Mieda-Sato, 2012). The loss of nuclear location signals in TDP-43 facilitates the cytoplasmic presentation and profoundly changes the miRNA profiles(Paez-Colasante *et al*, 2020). Interestingly, genetic manipulation of wild-type TDP-43 in motor neuron-like cells hinders miRNA biogenesis(Emde *et al*, 2015) and facilitates the formation of stress granules (SGs) that interact with and impede miRNA processing(Chen & Cohen, 2019). However, the specific mechanisms connecting TDP-43 pathological mutations with aberrant miRNA maturation remain uncertain, and clear evidence for either loss of normal function or gain of toxic function in existing models is lacking.

In this study, we aimed to explore the contribution of miRNA dicing to the pathogenic mutations of TDP-43-caused cytotoxicity and degeneration in vivo and in vitro. We found that null mutation in TDP-43 *Drosophila* homolog TBPH, as well as the counterparts A315T mutations, had a pronounced impact on Dicer activation and selectively mediated miRNA maturation in fly models. Manipulating Dicer expression differentially impacts A315T and M337V mutation-induced tissue degeneration and miRNA maturation, providing a distinctive molecular linkage to the disease mutation outputs. Furthermore, oxidative stress promotes TDP-43 A315T, and M337V redistributes to SG and leads to Dicer complex instability, but M337V preserves the functional Dicer activity in vitro. Our study expanded the understanding of the pathogenic mutations in TDP-43 influence miRNA dicing in ALS models, offering a basis for uncovering new mechanistic insights into the neurodegeneration and locomotor impairments induced by these mutations.

## Results and discussion

### TBPH deficiency downregulates DCR links to dysregulated miRNA and locomotor impairment in fly

To define potential pathways of miRNA processing involved in TDP-43-associated disease progression, we initially used null mutation of *Drosophila* TDP-43 (dTDP-43/TBPH)(Feiguin *et al*, 2009) to assess the relevant molecular expression. A significant decrease in the mRNA levels of *DCR-1*, *DCR-2*, *Drosha*, *Parsha*, *R2D2*, and *Loqs* was observed in flies carrying either TBPH^+/-^ or TBPH^-/-^ (Fig. EV1E). DCR-1, DCR-2, and Drosha showed reduced protein levels in TBPH^-/-^ flies, while R2D2 did not exhibit any changes in protein levels (Fig. 1A-D).

**Fig 1.**
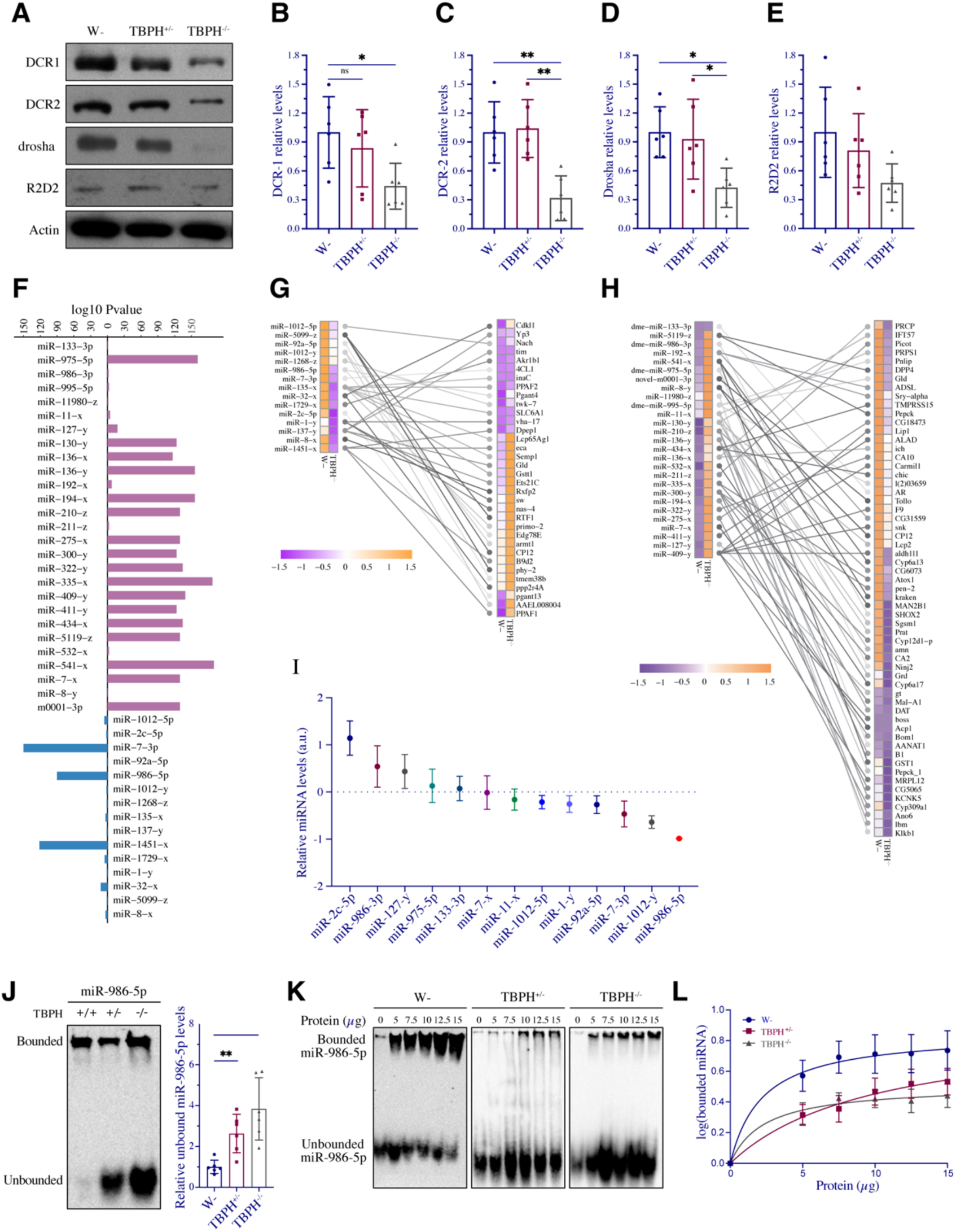
TBPH deficiency downregulates DCR levels and dysregulates miRNA processing. (**A-E**) Western blot analyses of the effect of DCR-1, DCR-2, Drosha, and R2D2 in TBPH^+/-^ and TBPH^-/-^ flies, and the relative protein expression were quantified (n=6 flies for each genotype) (**F**) miRNA-seq derived 42 most differentially expressed miRNAs. qPCR assay for comparing the mRNA levels of *DCR-1*, *DCR-2*, *Drosha*, *Pasha*, *R2D2,* and *Loqs* between W- and TBPH^-/-^ (n=6, biological replicates). Data were plotted as fold change from W-control. (**G**) Regulatory networks of downregulated miRNA and their targeted upregulated mRNA in TBPH^-/-^ flies. (**H**) Regulatory networks of upregulated miRNA and their targeted downregulated mRNA in TBPH^-/-^ flies. (**I**) RT-qPCR validation some of the differential expressed miRNAs altered in the TBPH^-/-^ flies (n=6 for each genotype). (**J**) EMSA and quantification analysis of the hybrid capacity of miR-986-5p with protein lysates from control (TBPH^+/+^), heterozygous (TBPH^+/-^), and homozygous TBPH (TBPH^-/-^). (**K**) The EMSA analysis for dme-miR-986-5p (1μM) binding with various amounts of protein lysates (dose plot range from 0 to 15 μg) from flies. (**L**) The density of bound signals in each group was measured and the results were analyzed using nonlinear regression by plotting the various amounts of protein. Data are shown as mean ± SEM and were analyzed using a one-way ANOVA with Tukey’s multiple comparisons test. *p < 0.05, **p < 0.01.

To understand the impact of TBPH on miRNA processing and potentially target and regulate mRNA expression, we carried out RNA-Seq and miRNA-Seq and integrated the analyzed miRNA and mRNA expressions. Through analysis of a two-fold change and a false discovery rate (FDR) of less than 0.05, we identified 336 genes that exhibited significant differential expression patterns between control and TBPH^-/-^ flies. Among these, 105 mRNAs were downregulated, and 201 mRNAs were upregulated. (Fig. EV1A-B and Datasets EV1). Meanwhile, 42 miRNAs were differentially expressed in the TBPH^-/-^ group, with 15 miRNAs showing reduced and 27 miRNAs showing upregulated (Fig. EV1C-D and Datasets EV2). Under the integrated analysis, the downregulated miRNAs (15) targeted 34 upregulated mRNAs, whereas the upregulated miRNAs (27) targeted 59 downregulated mRNAs have been correlated (Fig. 1F-H). Validating the levels of miRNAs and mRNAs using RT-qPCR analysis indicated that most of the dysregulated miRNAs were consistent with the miRNA-seq and mRNA-seq results (Fig. 1I and EV1F). Among them, miR-986-5p was the most significantly downregulated in TBPH-/- compared to the control (Fig. 1I and EV1D).

Previous studies have indicated that TDP-43 physically interacts with miRNAs and facilitates their processing(Kawahara & Mieda-Sato, 2012; Paez-Colasante *et al*., 2020). To gain insight into the effect of TBPH on miRNA binding affinity, we employed a native gel mobility shift assay to evaluate miR-986-5p, miR-7-5p, and miR-975-5p, which were drastically altered in TBPH null mutations. Quantitative analysis suggested that a significant amount of free miRNA in biotin-labeled miR-986-5p was incubated with proteins isolated from TBPH^+/-^ and TBPH^-/-^ flies (Fig. 1J). However, the unbound free bands of miR-7-5p and miR- 975-5p with the protein lysate of TBPH^+/-^ and TBPH^-/-^ showed no significant difference between the groups (Figs. EV1G, J).

We further evaluated the affinity profile of the TBPH effect on miRNA binding, and we measured varying amounts of protein affinity with biotinylated probes. The results indicated that with increasing amounts of protein lysates from W-, the bands of the protein-miRNA complex increased, but free miR-986-5p gradually decreased. The apparent binding affinities (EC50s) of the W-, TBPH^+/-^, and TBPH^-/-^ groups for probe binding were determined using the titration method(Paez-Colasante *et al*., 2020). For miR-986-5p, W- had the highest apparent affinity (EC50 = 3.651) and a significantly lower binding affinity for TBPH^+/-^ (EC50 = 4.114) and TBPH^-/-^ (EC50 = 4.329) (Figs. 1K, L). The apparent affinity of miR-7-5p was observed in W- (EC50 = 2.868), TBPH^+/-^ (EC50 = 2 .589), and TBPH^-/-^ (EC50 = 1.922) (Figs. EV1H, I).

Similarly, for miR-975-5p, an apparent affinity was found in W- (EC50 = 2.263), TBPH^+/-^ (EC50 = 2.099), and TBPH^-/-^ (EC50 = 1.922) (Figs. EV1K, L). Taken together, these results suggest that TBPH preferentially affects the binding affinity of miR-986-5p.

Although we demonstrated a close association between miR-986-5p and TBPH in flies through in vitro binding experiments, it is unclear whether miR-986-5p contributes to TBPH- related phenotypic impairments. We tested the spontaneous motility of flies with miR-986 knockout or overexpression without one copy of TBPH(Long *et al*, 2023). Similar to the TBPH^+/-^, heterogeneity of miR-986-5p knockout (miR-986-5p^+/-^) resulted in impairment of movement distance and velocity compared to W- (Fig. EV2B-D). However, ubiquitous expression of miR-986-5p in TBPH^+/-^ flies effectively restored the motility defects to a normal level, even though the miR-986-5p levels reached approximately half of the W- group (Fig. EV2A). The findings indicate that TBPH-restricted exogenous miR-986-5p expression concurrently exhibits a restorative function of miR-986-5p on the locomotor impairment in flies with TBPH deficiency. We confirmed that expression miR-986-5p or its sponge in the *Drosophila* compound eye did not influence ommatidia damage (Fig. EV2F).

To further investigate these possibilities, we administered small molecular enoxacin to TBPH-null flies, which has been reported to facilitate Dicer recruits to AGO2 complex that enhance miRNA and siRNA biogenesis(Shan *et al*, 2008; Zhang *et al*, 2008). To elucidate the binding pattern of enoxacin to *Drosophila* Dicer, we initially crystallized Dicer-1 and Dicer-2 and subjected to ligand soaking to obtain complex structures with enoxacin (1.5 Å resolution) (Fig. EV2E). Enoxacin shares a binding mode with a pocket formed by the RNase IIIa/b domains of DCR-1 and DCR-2 proteins, respectively (Fig. EV2E, Table EV1). Following treatment with enoxacin, TBPH^+/-^ larvae exhibited increased movement, turning, and forward movements, whereas TBPH^-/-^ larvae showed an increase in reverse movement to a certain extent (Fig. EV1G). In our results, TBPH deficiency causes miR-986 processing bias to reduce mature 5p strand but elevate 3p strand expression. Enoxacin treatment profoundly promotes miR-986- 5p and its counterpart 3p strand expression. However, TBPH deficiency restricts the increase of miR-986-5p but not the 3p strand (Fig. EV2H, I). Together with the observation that TBPH restricts exogenous miR-986-5p expression in fly tissue, these findings prove that TBPH modulates miRNA expression by suppressing the function of the Dicer protein.

### Requirements of DCR-1 in TBPH A315T-induced cytotoxicity and degenerative outputs

Although TBPH is crucial for Dicer processing miRNAs, which are accompanied by significant motor impairment, it remains unclear whether disease-related mutations influence DCRs and/or miRNA alterations. We investigated the genetic interactions between Dicer proteins and wild-type (WT) TBPH or an A315T mutant, corresponding to a missense mutation found in humans (Fig. EV3A). To evaluate the possible influence of TBPH A315T on DCRs, we investigated the expression levels of DCR-1, DCR-2, Drosha, and R2D2 in adult flies harboring wild-type (WT) and A315T mutations in TBPH. The WB analysis demonstrated that both TBPH WT and TBPH A315T mutations led to a significant reduction in the levels of DCR- 1 and DCR-2 compared to the LacZ control, with the A315T mutation demonstrating a more pronounced reduction (Figs. 2A-C). However, no significant difference was observed in Drosha and R2D2 levels between the groups (Figs. 2D, E). Since DCR-1 is indispensable for endogenous miRNA biogenesis, therefore we focused on the investigation of the interaction of TBPH A315T with DCR-1 in flies.

**Fig 2.**
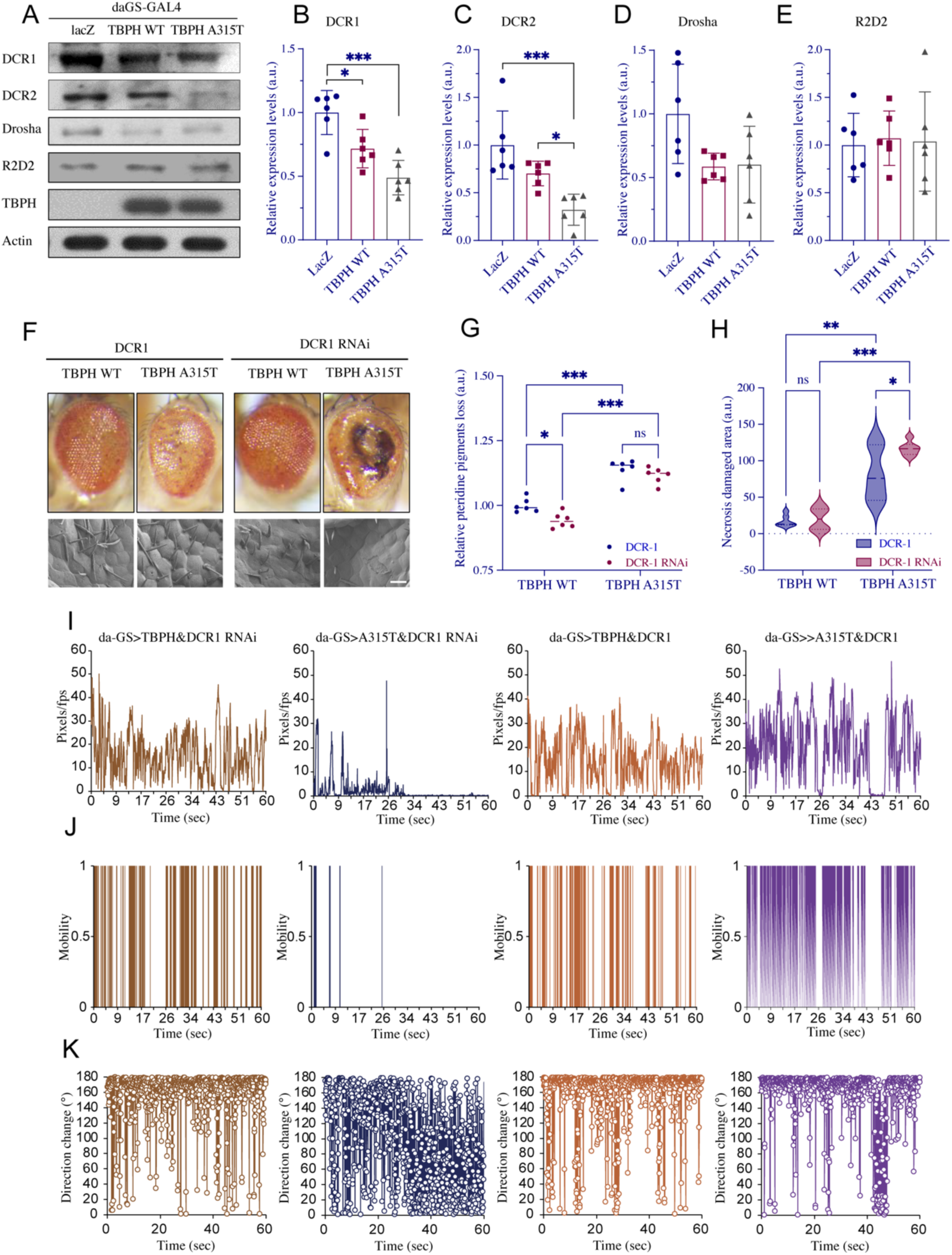
The genetic interaction between DCR-1 and the TBPH A315T mutation impacts compound eye degeneration and locomotor behavior. (**A**) Western blot analysis of DCR-1, DCR-2, Drosha, and R2D2 levels in daGS-Gal4>LacZ (lane 1), daGS-Gal4>TBPH WT (lane 2), and daGS-Gal4>TBPH A315T (lane 3) induced by 200 μM RU486. The blots were plotted using actin as a loading control. The relative levels of DCR1 (**B**), DCR2 (**C**), Drosha (**D**), and R2D2 (**E**) were quantified independently. (n=3, biological replicates). (**F**) Light (upper) and scanning electron microscopy (lower) images of fly eyes (5-day old) show that RNAi or DCR1 overexpression affects retinal degeneration in flies expressing the TBPH A315T mutation under the control of the GMR-Gal4 driver. Scale bars, 15 μm. Quantification of damaged areas (**G**) and pteridine pigmentation loss (**F**) in fly eyes with DCR1 RNAi or DCR1 overexpression in TBPH WT or TBPH A315T on day 5. (n=6, for each genotype). (**I**-**K**) Knockdown of DCR1 expression by RNAi attenuates DCR1 RNAi, DCR1 overexpression, and spontaneous locomotion impairment in TBPH A315T mutant flies. Representative images of spontaneous locomotion with parameters of movement (**I**), immobility (**J**), and direction changes (**K**) with a 1 min plot. Data were analyzed using one-way ANOVA with Tukey’s post-hoc test. (*p < 0.05, **p < 0.001; ns, not significant). Error bars represent mean ± SEM.

We first evaluated the expression of TBPH WT and TBPH A315T impacts on compound eye degeneration. Consistent with the report(Estes *et al*, 2011), expression of TBPH WT and TBPH A315T in fly eyes (Gmr-Gal4>) leads to adult lethality in males and females under a 25℃ environment. Therefore, we dissected the pupa and found reduced pigmentation in TBPH WT and TBPH A315T flies. In addition, the compound eye of TBPH A315T flies also has black necrotic damage in both male and female pupae (Fig. EV3B). The fly larvae were then maintained at 18℃ to appropriately mitigate the toxicity of gene expression and obtain successful birth production. Based on previous reports, we quantified this prominent eye damage by analyzing pteridine pigment loss(Mitra & Ryoo, 2021; Ooi *et al*, 1997) and necrosis- damaged area(Appocher *et al*, 2017). At lower temperatures, flies expressing TBPH WT and TBPH A315T still showed reduced eye pigmentation, which deepened gradually with age (Fig. EV3C). We observed that the A315T flies had more pteridine pigment loss (Fig. EV3D) and a relatively larger area of necrotic damage compared to TBPH WT flies on day 5 (Fig. EV3E).

To investigate the potential impact of Dcr on the pathogenic effects of TBPH, we manipulated DCR-1 or DCR-1 RNAi with the co-expression of TBPH WT and A315T mutant, specifically in the compound eyes of flies. We subsequently examined the alterations in eye degeneration in flies that DCR-1 or DCR-1 RNAi was co-expressed with TBPH WT and A315T, respectively. The ommatidia organization in TBPH A315T has a significantly reduced pteridine pigment and increased necrosis-damaged area compared to TBPH WT (Figs. EV3C-E). (Fig. EV3F). Intriguingly, DCR-1 RNAi reduced pteridine pigmentation loss in TBPH WT but not TBPH A315T compound eyes (Figs. 2F-G). DCR-1 RNAi or expression did not change the relative necrotic damage area in TBPH WT flies. However, DCR-1 RNAi significantly induced retinal damage in TBPH A315T flies (Figs. 2F, H).

Subsequently, locomotor behavior was assessed using daGS-Gal4 to induce the expression of TBPH WT and TBPH A315T in flies under RU486 administration(Robles-Murguia *et al*, 2019). Without RU486 induction, the flies in each group exhibited normal movement ability.

Under RU486 induction, TBPH WT and TBPH A315T expression exhibited a pronounced reduction in locomotor activity compared to age-matched LacZ (Figs. EV4 A-C). TBPH WT and TBPH A315T flies showed drastic impairments in locomotor behavior, including reduced movement distance (Fig. EV4B) and increased immobility time (Fig. EV4C). Notably, TBPH A315T flies displayed a more severe motor impairment than the TBPH WT flies in response to RU486 at 0 µM, 10 µM, 100 µM, and 200 µM concentrations. (Figs. EV4B, C).

Since there was no significant drug-concentration-dependent impairment in motor ability among the groups, we used a small concentration of 10 μM RU486 to induce the expression of TBPH WT and TBPH A315T for subsequent experiments. TBPH WT and TBPH A315T expression regulated the majority of the dysregulated miRNAs identified in TBPH^-/-^ flies (Fig. EV4E), with a pronounced severe disrupted affinity binding (Fig. EV4 F) compared to TBPH WT. Consistently, DCR-1 RNAi significantly reduced the movement distance (Fig. 2I), mobility status (Fig. 2J), and disrupted direction change (Fig. 2K) in TBPH A315T compared to TBPH WT flies, the expression of DCR-1 improved locomotion. In the motor neuron expression (D42-Gal4>) of TBPH WT and A315T, the proteins colocalized with endogenous DCR-1 and DCR-2 (Fig. EV4D). Together, these results indicate that the TBPH A315T disrupted the cytoplasm Dicer-1 functionally processing miRNA.

### TDP-43 mutations differentially regulating DCR that related to compound eye degeneration

To determine whether mutations of human TDP-43 have the function of regulating Dcr expression in flies, we examined WT, A315T, and M337V mutations in TDP-43 expressed in the fly thorax. WB results revealed a slight decrease in the expression levels of DCR-1 and DCR-2 in TDP-43 WT. Unexpectedly, the TDP-43 A315T mutant exhibited a substantial increase in DCR-2 expression, but it did not affect DCR-1 levels. However, the TDP-43 M337V mutant showed an increase in DCR-1 expression compared to LacZ and an increase in DCR-2 expression compared to WT (Figs. 3A-C). We subsequently assessed the retinal organization of the compound eyes by expressing TDP-43 WT, TDP-43 A315T, or TDP-43 M337V. TDP-43 A315T and TDP-43 M337V expression in flies resulted in a significant loss of pteridine pigments in compound eyes compared to the TDP-43 WT. The loss of compound eye pteridine pigments is more severe in TDP-43 A315T mutations compared to M337V mutations (Figs. EV4G, H). Next, we explored whether RNAi or the genetic expression of DCR-1 or DCR-2 could affect compound eye degeneration in flies expressing TDP-43 WT, the A315T, and M337V pathological mutations. The expression of DCR-1 had a slight suppressive effect on the loss of pteridine pigment in TDP-43 WT flies. However, DCR-1 RNAi significantly promoted the loss of red pteridine pigment (Figs. 3D, E). In addition, knockdown or expression of DCR- 1 and DCR-2 did not alter the loss of pteridine pigment in TDP-43 A315T mutants. Intriguingly, changing the expression of DCR-1 profoundly affected the TDP-43 M337V mutation, with DCR-1 expression counteracting compound eye degeneration, while DCR-1 RNAi ameliorated eye degeneration (Fig. 3E). Altering DCR-2 expression had a minor impact on compound eye degeneration caused by the TDP-43 A315T and M337V mutations.

**Fig 3.**
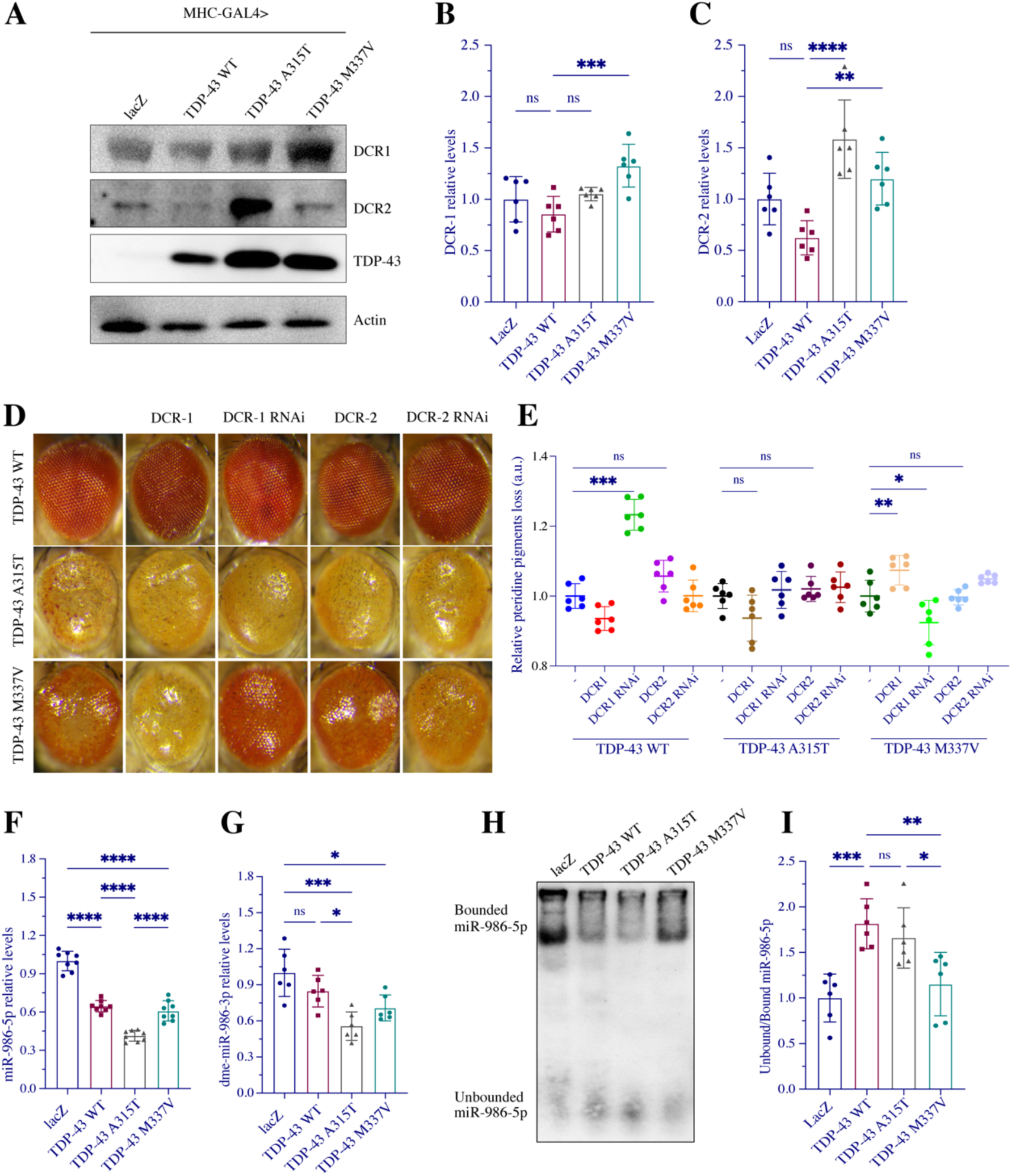
TDP-43 pathological mutations differentially interact with DCR-1 and DCR-2 and affect miRNA binding affinity. (**A**) Western blot analysis of DCR1, DCR2 levels in LacZ (lane 1), TDP-43 WT (lane 2), TDP-43 A315T (lane 3), and TDP-43 M337V (lane 4) under the control of MHC-Gal4. Quantified protein expression of DCR1 (**B**) and DCR2 (**C**) in individual groups. (**D**) Light microscopy images of eye defects from 5-day-old flies with Gmr-Gal4 derived TDP-43 WT, TDP-43 A315T, and TDP-43 M337V. (**E**) Quantification of the degenerated area (relative pteridine pigment loss) in fly eyes expressing knockdown or overexpression of DCR-1 and DCR-2 in the indicated groups of wild-type (WT), A315T-TDP-43, or M337V-TDP-43. (**F**) RT-qPCR to detect dme-miR-986-5p and its *Star 3p chain in flies with wild-type, A315T, and M337V of TDP-43 mutations (n=6, biological replicates). (**G**) EMSA analysis of the unbounded dme-miR-986-5p with lysate of flies expressing TDP-43 WT, TDP-43 A315T, or TDP-43 M337V under the control of the MHC-Gal4 driver. MHC-Gal4>LacZ as control. (**H**) EMSA analysis of the binding affinity miR-986-5p with protein lysates from fly-expressed TDP-43 WT, TDP-43 A315T, and TDP-43 M337V mutations. (**I**) Quantifying the binding efficiency of miR-986-5p with protein lysate from fly-expressed TDP-43 WT, TDP-43 A315T, and TDP-43 M337V mutations. (n=6, biological replicates). Data were analyzed using one-way ANOVA with Tukey’s post hoc test (*p < 0.05, **p < 0.01, ***p < 0.0001). Error bars represent mean ± SEM.

Since the Dcr proteins interacted specifically with the TDP-43 WT, A315T, and M337V mutations, we speculated that the levels of endogenous miRNAs, such as miR-986, might be altered in flies harboring them. The TDP-43 WT, A315T, and M337V mutations decreased the levels of miR-986-5p and its counterpart chain miR-986-3p compared to the LacZ control, with the A315T mutation having the most pronounced effect (Figs. 3F, G). The result is reminiscent of that ectopic expression of TDP-43 M337V produces a less severe sensory organ precursor (SOP) phenotype, which is comparatively less severe than that observed in TDP-43 WT and A315T in fly(Li *et al*, 2013) and mouse models(Wang *et al*, 2017). The supporting evidence has been obtained in EMSA that the binding affinity of miR-986-5p to the proteins was significantly reduced by both the WT and A315T mutations of TDP-43, whereas the M337V mutation of TDP-43 had no significant impact on miRNA-protein binding (Figs. 3H, I). The findings indicate that in contrast to TDP-43 WT, pathological mutations in TDP-43 govern miRNA biogenesis by altering endogenous DCR homologous interactions in distinct ways.

### Pathogenic mutations redistribute TDP-43 and differentially vulnerable Dicer activity

It has been proposed that the interaction between multiple RNA-binding proteins, including TDP-43, typically located in stress granules (SGs), and either Dicer or Ago2(Emde *et al*., 2015; Kawahara & Mieda-Sato, 2012; Li *et al*, 2012). Mutations in TDP-43 in ALS have been found to trigger a stress response that leads to the formation of SG. To determine the potential pathways underlying TDP-43 mutations induced disease progression, we transfected TDP-43 WT and pathological mutations and evaluated its localization to formed aggregates using specific markers under an oxidative condition(Emde *et al*., 2015). Cells transfected with TDP-43 mutations promoted its cellular colocalization with TIAR (SGs marker) (Figs. EV5C- E), Dicer (Figs. 4A, C and D), and AGO2 (Figs. 4G-I) compared to TDP-43 WT and empty control. For example, approximately 25% of the cells expressing TDP-43 displayed TDP-43- Dicer aggregates; a higher percentage (approximately 60%) was observed in cells transfected with TDP-43 A315T and M337V mutations (Fig. 4A, C). TDP-43 WT, A315T, and M337V mutations transfection did not change Dicer (Fig. 4D), Drosha, AGO1, and AGO2 expression drastically (Fig. EV4I). However, H_2_O_2_ treatment reduced Dicer levels in the TDP-43 mutation- transfected cells compared to WT and control cells (Fig. 4E, F), while levels of Drosha, AGO1, and AGO2 were unaffected (Fig. EV4K). Quantification of TDP-43-Dicer coaggregation under 400 μM H_2_O_2_ condition suggested that different to WT, the presence of aggregate positive cells transfected in TDP-43 A315T and M337V was restricted compared to the WT group (Fig. 4A-C). Meanwhile, oxidative treatment elevated TDP-43 colocalizing with Dicer in the empty, TDP-43 WT, and A315T transfection groups, whereas M337V transfection restricted the formation of aggregates (Fig. 4A, B, and D). In addition, the TDP-43 A315T and M337V mutations reduced the formation of aggregates with AGO2, the key component of RISCs, under oxidative conditions (Fig. 4G, J). These observations suggest that TDP-43 mutations result in the redistribution and aggregation of proteins, potentially exhibiting a preference for recruiting the Dicer complex to AGO2.

**Fig 4.**
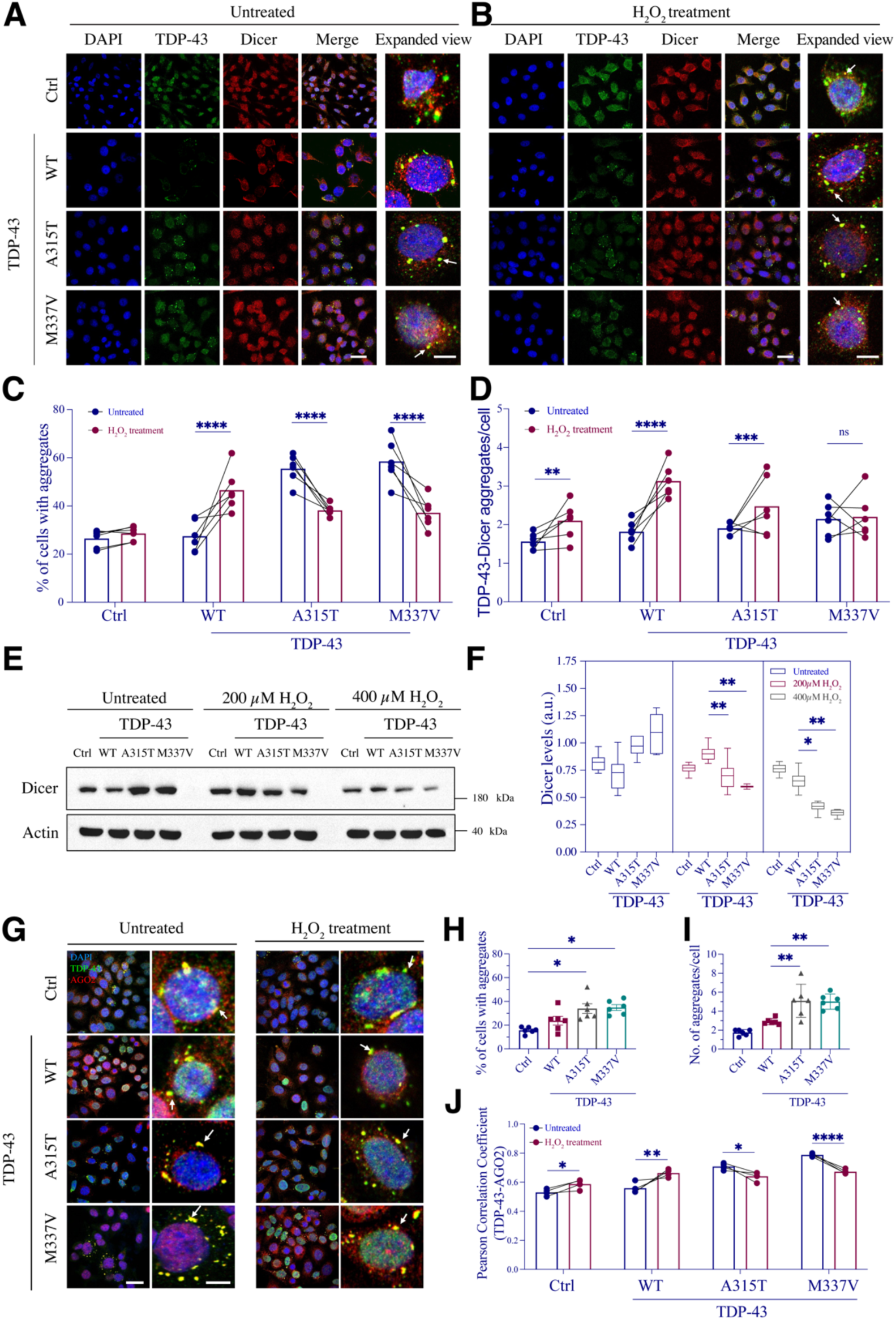
TDP-43 pathological mutations disrupt the interaction with Dicer. (**A**-**B**) HeLa cells were transfected with empty control, TDP-43 wild-type (WT), A315T, and M337V mutants for 36 hours and then were treated without (**A**) or with 400 μM H_2_O_2_ for four hours (**B**). Scale bars, 20 μm (left); 5 μm (right). (**C**) Comparison of the proportion (%) of cells with TDP-43-Dicer aggregates in (**A**) and (**B**) conditions. % of positive cells = number of cells with TDP-43-Dicer aggregates/total number of cells in selected area x 100%. (n=6, biological replicates). (**D**) Comparison of the changing TDP-43-Dicer aggregates within the cells under untreated (**A**) or oxidative treatment (**B**) conditions. (**E**) HeLa cells transfected with wild-type, A315T, and M337V mutations of TDP-43 for 36 hours and then treated with H_2_O_2_ in a dose of 0 μM, 200 μM, or 400 μM for four hours, respectively. Whole cells were lysed and analyzed by western blot with the Dicer antibody. Actin was used as the loading control. (**F**) The Dicer protein levels were quantified and normalized relative to actin. (n=6, biological replicates). (**G**) Following transfection, HeLa cells were treated with 400 μM H_2_O_2_ for four hours and then immunofluorescence (IF) staining with anti-AGO2 (red) and TDP-43 (green) antibodies. Nuclei were counterstained with DAPI (blue) in merged images. The cells were then visualized by confocal microscopy. % of cells (**H**) with TDP-43-AGO2 aggregates and the number of double positive aggregates per cell (**I**) were quantified (n=6, biological replicates). (**J**) Pearson’s correlation coefficients between TDP-43 and AGO2 on the results shown in (**G**). Data were analyzed using a two-tailed Student’s *t*-test or one-way ANOVA with Tukey’s post-hoc test (*p < 0.05, **p < 0.01, ***p < 0.0001, ****p < 0.00001, ns, not significant). Error bars represent mean ± SEM.

Dicer proteins have a structured helicase domain on their N-terminus, followed by DUF283 and PAZ domains, and tandem RNase III domains at their C-terminus (Fig. EV5A). Consisting with *Drosophila* DCR-1 and DCR-2, human Dicer shares a similar binding mode, with a pocket formed by the RNase IIIa/b domain contact with enoxacin (Fig. EV5D). The protein architecture of humans and *Drosophila* Dicer resembles an L-shaped inactive form, with an opening in the helicase domain for loading miRNA substrates that subsequently convert into the active form(Zapletal *et al*, 2023; Zapletal *et al*, 2022). Duplex short RNAs (miRNAs or siRNAs) derived from Dicer catalytic activity in the cytosol are essential for translational repression and degradation of targeted mRNA(Leung *et al*, 2006). We conducted a quantitative analysis of three miRNAs: hs-miR-143, hs-miR-558, and hs-miR-574, known to be directedly bound to human TDP-43(Kawahara & Mieda-Sato, 2012) with or without enoxacin treatment. Our results demonstrated relatively reduced levels of hs-miR-574 in TDP-43 WT and A315T mutation, even though the decrease did not get a significant difference in hs-miR-143 and hs- miR-558 (Figs. EV5G-I). However, enoxacin significantly elevated the hs-miR-558 and hs- miR-574 expression in M337V mutation cells compared to TDP-43 A315T (Fig. EV5I). The enoxacin treatment did not alter the Dicer and AGO2 levels (Fig. EV5F).

These findings indicate that conditional stress redistributes TDP-43 to form cellular SGs and localize to RISCs, and TDP-43 mutations restrict interaction with Dicer proteins and lead to Dicer complex instability. Together with the observation in *Drosophila*, the TDP-43 A315T mutation disrupts the interaction with and functional integration of Dicer. The M337V mutation hinders the interaction with Dicer as well, while the functional integrity of Dicer, such as their catalytic activity, is still preserved to some extent.

## Acknowledgments

We thank Nobutaka Hattori and Yuzuru Imai at the Department of Neurology, Juntendo University, Tokyo, Japan, for their kind gift of fly lines and helpful suggestions and discussions regarding the *Drosophila* experiments. We also thank Professor C.-K. James Shen, Institute of Molecular Biology, Academia Sinica, Taipei, Taiwan, for sharing the plasmids. We also thank Professor Fabian Feiguin, International Centre for Genetic Engineering and Biotechnology, Trieste, Italy, for the gift of TBPH null fly lines. We thank the Bloomington Fly Stock Center (BDRC) and the Center for Excellence in Molecular Cell Science at the CAS (Shanghai, China) for providing the fly lines. This work was supported by the National Natural Science Foundation of China (NSFC) (82171414), Suzhou Science and Technology Plan Project (2022SS02), Natural Science Foundation of Jiangsu Province (BK20200192), and a special launch fund from Soochow University.

## Author contributions

H. R. M. and X. L. conceptualized and designed the study, and wrote the first version of the manuscript. Sample preparation, data collection, and analyses were performed by X. L., M. N. J., H. H. D., and Y. X. M.. L. L. C. and J. L. performed methodology and data validation. C. F. L., H. R. M., and J. L. acquired funding for this project. C. F. L. and H. R. M. wrote and edited the final version of the manuscript and supervised the study. All the authors commented on, approved, and agreed to the published version of the manuscript.

## Materials and Methods

### Reagents and Tools Table

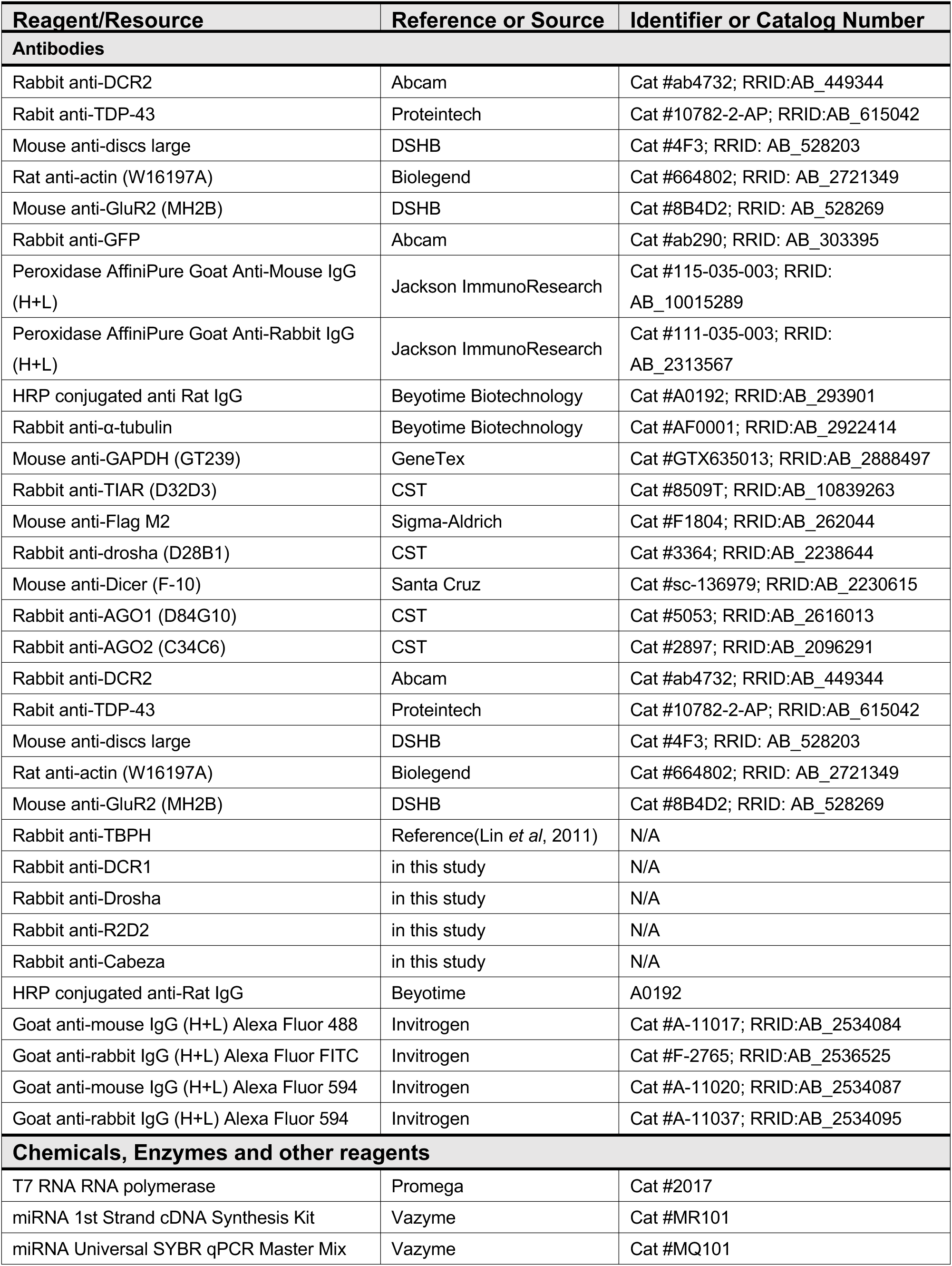
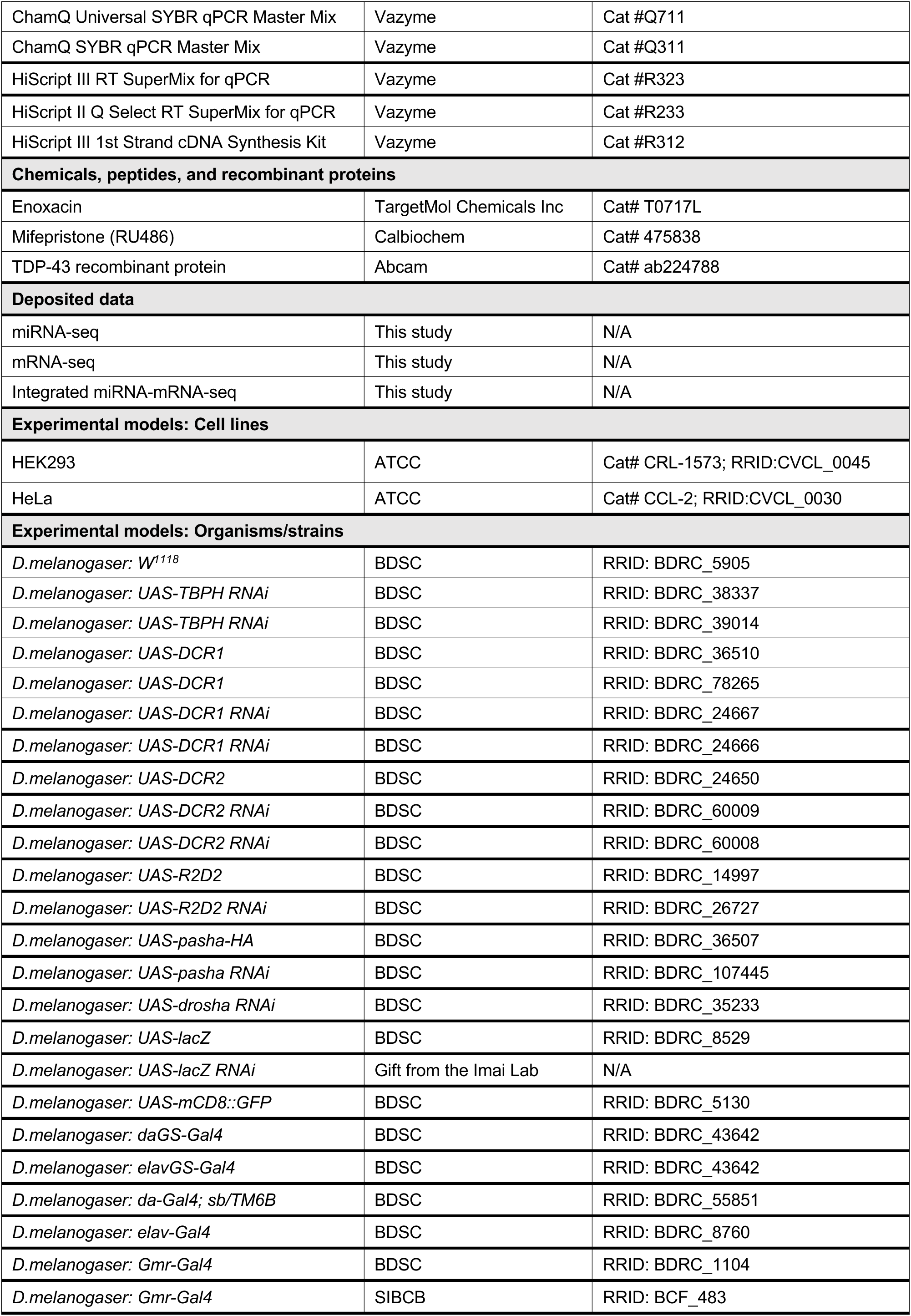
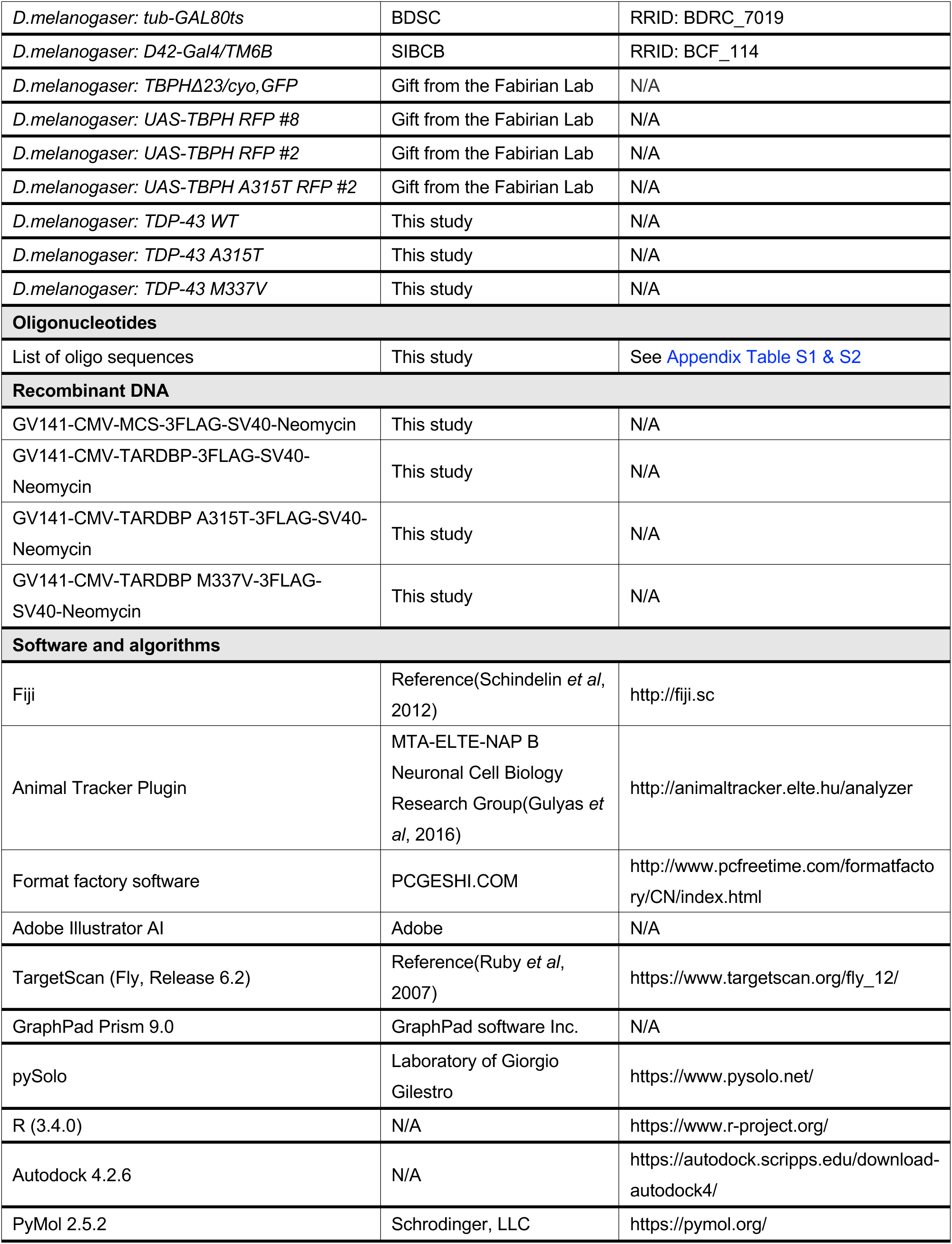

| Reagent/Resource | Reference or Source | Identifier or Catalog Number |
| --- | --- | --- |
| **Antibodies** |  |  |
| Rabbit anti-DCR2 | Abcam | Cat #ab4732; RRID:AB_449344 |
| Rabbit anti-TDP-43 | Proteintech | Cat #10782-2-AP; RRID:AB_615042 |
| Mouse anti-discs large | DSHB | Cat #4F3; RRID: AB_528203 |
| Rat anti-actin (W16197A) | Biolegend | Cat #664802; RRID: AB_2721349 |
| Mouse anti-GluR2 (MH2B) | DSHB | Cat #8B4D2; RRID: AB_528269 |
| Rabbit anti-GFP | Abcam | Cat #ab290; RRID: AB_303395 |
| Peroxidase AffiniPure Goat Anti-Mouse IgG (H+L) | Jackson ImmunoResearch | Cat #115-035-003; RRID: AB_10015289 |
| Peroxidase AffiniPure Goat Anti-Rabbit IgG (H+L) | Jackson ImmunoResearch | Cat #111-035-003; RRID: AB_2313567 |
| HRP conjugated anti Rat IgG | Beyotime Biotechnology | Cat #A0192; RRID:AB_293901 |
| Rabbit anti-α-tubulin | Beyotime Biotechnology | Cat #AF0001; RRID:AB_2922414 |
| Mouse anti-GAPDH (GT239) | GeneTex | Cat #GTX635013; RRID:AB_2888497 |
| Rabbit anti-TIAR (D32D3) | CST | Cat #8509T; RRID:AB_10839263 |
| Mouse anti-Flag M2 | Sigma-Aldrich | Cat #F1804; RRID:AB_262044 |
| Rabbit anti-drosha (D28B1) | CST | Cat #3364; RRID:AB_2238644 |
| Mouse anti-Dicer (F-10) | Santa Cruz | Cat #sc-136979; RRID:AB_2230615 |
| Rabbit anti-AGO1 (D84G10) | CST | Cat #5053; RRID:AB_2616013 |
| Rabbit anti-AGO2 (C34C6) | CST | Cat #2897; RRID:AB_2096291 |
| Rabbit anti-DCR2 | Abcam | Cat #ab4732; RRID:AB_449344 |
| Rabbit anti-TDP-43 | Proteintech | Cat #10782-2-AP; RRID:AB_615042 |
| Mouse anti-discs large | DSHB | Cat #4F3; RRID: AB_528203 |
| Rat anti-actin (W16197A) | Biolegend | Cat #664802; RRID: AB_2721349 |
| Mouse anti-GluR2 (MH2B) | DSHB | Cat #8B4D2; RRID: AB_528269 |
| Rabbit anti-TBPH | Reference(Lin *et al*, 2011) | N/A |
| Rabbit anti-DCR1 | in this study | N/A |
| Rabbit anti-Drosha | in this study | N/A |
| Rabbit anti-R2D2 | in this study | N/A |
| Rabbit anti-Cabeza | in this study | N/A |
| HRP conjugated anti-Rat IgG | Beyotime | A0192 |
| Goat anti-mouse IgG (H+L) Alexa Fluor 488 | Invitrogen | Cat #A-11017; RRID:AB_2534084 |
| Goat anti-rabbit IgG (H+L) Alexa Fluor FITC | Invitrogen | Cat #F-2765; RRID:AB_2536525 |
| Goat anti-mouse IgG (H+L) Alexa Fluor 594 | Invitrogen | Cat #A-11020; RRID:AB_2534087 |
| Goat anti-rabbit IgG (H+L) Alexa Fluor 594 | Invitrogen | Cat #A-11037; RRID:AB_2534095 |
| **Chemicals, Enzymes and other reagents** |  |  |
| T7 RNA RNA polymerase | Promega | Cat #2017 |
| miRNA 1st Strand cDNA Synthesis Kit | Vazyme | Cat #MR101 |
| miRNA Universal SYBR qPCR Master Mix | Vazyme | Cat #MQ101 |
| ChamQ Universal SYBR qPCR Master Mix | Vazyme | Cat #Q711 |
| ChamQ SYBR qPCR Master Mix | Vazyme | Cat #Q311 |
| HiScript III RT SuperMix for qPCR | Vazyme | Cat #R323 |
| HiScript II Q Select RT SuperMix for qPCR | Vazyme | Cat #R233 |
| HiScript III 1st Strand cDNA Synthesis Kit | Vazyme | Cat #R312 |
| **Chemicals, peptides, and recombinant proteins** |  |  |
| Enoxacin | TargetMol Chemicals Inc | Cat# T0717L |
| Mifepristone (RU486) | Calbiochem | Cat# 475838 |
| TDP-43 recombinant protein | Abcam | Cat# ab224788 |
| **Deposited data** |  |  |
| miRNA-seq | This study | N/A |
| mRNA-seq | This study | N/A |
| Integrated miRNA-mRNA-seq | This study | N/A |
| **Experimental models: Cell lines** |  |  |
| HEK293 | ATCC | Cat# CRL-1573; RRID:CVCL_0045 |
| HeLa | ATCC | Cat# CCL-2; RRID:CVCL_0030 |
| **Experimental models: Organisms/strains** |  |  |
| *D.melanogaster*: *W<sup>1118</sup>* | BDSC | RRID: BDRC_5905 |
| *D.melanogaster*: *UAS-TBPH RNAi* | BDSC | RRID: BDRC_38337 |
| *D.melanogaster*: *UAS-TBPH RNAi* | BDSC | RRID: BDRC_39014 |
| *D.melanogaster*: *UAS-DCR1* | BDSC | RRID: BDRC_36510 |
| *D.melanogaster*: *UAS-DCR1* | BDSC | RRID: BDRC_78265 |
| *D.melanogaster*: *UAS-DCR1 RNAi* | BDSC | RRID: BDRC_24667 |
| *D.melanogaster*: *UAS-DCR1 RNAi* | BDSC | RRID: BDRC_24666 |
| *D.melanogaster*: *UAS-DCR2* | BDSC | RRID: BDRC_24650 |
| *D.melanogaster*: *UAS-DCR2 RNAi* | BDSC | RRID: BDRC_60009 |
| *D.melanogaster*: *UAS-DCR2 RNAi* | BDSC | RRID: BDRC_60008 |
| *D.melanogaster*: *UAS-R2D2* | BDSC | RRID: BDRC_14997 |
| *D.melanogaster*: *UAS-R2D2 RNAi* | BDSC | RRID: BDRC_26727 |
| *D.melanogaster*: *UAS-pasha-HA* | BDSC | RRID: BDRC_36507 |
| *D.melanogaster*: *UAS-pasha RNAi* | BDSC | RRID: BDRC_107445 |
| *D.melanogaster*: *UAS-drosha RNAi* | BDSC | RRID: BDRC_35233 |
| *D.melanogaster*: *UAS-lacZ* | BDSC | RRID: BDRC_8529 |
| *D.melanogaster*: *UAS-lacZ RNAi* | Gift from the Imai Lab | N/A |
| *D.melanogaster*: *UAS-mCD8::GFP* | BDSC | RRID: BDRC_5130 |
| *D.melanogaster*: *daGS-Gal4* | BDSC | RRID: BDRC_43642 |
| *D.melanogaster*: *elavGS-Gal4* | BDSC | RRID: BDRC_43642 |
| *D.melanogaster*: *da-Gal4; sb/TM6B* | BDSC | RRID: BDRC_55851 |
| *D.melanogaster*: *elav-Gal4* | BDSC | RRID: BDRC_8760 |
| *D.melanogaster*: *Gmr-Gal4* | BDSC | RRID: BDRC_1104 |
| *D.melanogaster*: *Gmr-Gal4* | SIBCB | RRID: BCF_483 |
| *D.melanogaster: tub-GAL80ts* | BDSC | RRID: BDRC_7019 |
| *D.melanogaster: D42-Gal4/TM6B* | SIBCB | RRID: BCF_114 |
| *D.melanogaster: TBPHΔ23/cyo, GFP* | Gift from the Fabirian Lab | N/A |
| *D.melanogaster: UAS-TBPH RFP #8* | Gift from the Fabirian Lab | N/A |
| *D.melanogaster: UAS-TBPH RFP #2* | Gift from the Fabirian Lab | N/A |
| *D.melanogaster: UAS-TBPH A315T RFP #2* | Gift from the Fabirian Lab | N/A |
| *D.melanogaster: TDP-43 WT* | This study | N/A |
| *D.melanogaster: TDP-43 A315T* | This study | N/A |
| *D.melanogaster: TDP-43 M337V* | This study | N/A |
| **Oligonucleotides** |  |  |
| List of oligo sequences | This study | See Appendix Table S1 & S2 |
| **Recombinant DNA** |  |  |
| GV141-CMV-MCS-3FLAG-SV40-Neomycin | This study | N/A |
| GV141-CMV-TARDBP-3FLAG-SV40-Neomycin | This study | N/A |
| GV141-CMV-TARDBP A315T-3FLAG-SV40-Neomycin | This study | N/A |
| GV141-CMV-TARDBP M337V-3FLAG-SV40-Neomycin | This study | N/A |
| **Software and algorithms** |  |  |
| Fiji | Reference(Schindelin *et al*, 2012) | http://fiji.sc |
| Animal Tracker Plugin | MTA-ELTE-NAP B<br>Neuronal Cell Biology<br>Research Group(Gulyas *et al*, 2016) | http://animaltracker.elte.hu/analyzer |
| Format factory software | PCGESHI.COM | http://www.pcfreetime.com/formatfactory/CN/index.html |
| Adobe Illustrator AI | Adobe | N/A |
| TargetScan (Fly, Release 6.2) | Reference(Ruby *et al*, 2007) | https://www.targetscan.org/fly_12/ |
| GraphPad Prism 9.0 | GraphPad software Inc. | N/A |
| pySolo | Laboratory of Giorgio Gilestro | https://www.pysolo.net/ |
| R (3.4.0) | N/A | https://www.r-project.org/ |
| Autodock 4.2.6 | N/A | https://autodock.scripps.edu/download-autodock4/ |
| PyMol 2.5.2 | Schrodinger, LLC | https://pymol.org/ |

## RESOURCE AVAILABILITY

### Lead contact

Further information and requests for resources and reagents should be directed to and will be fulfilled by the lead contact, Hongrui Meng.

### Methods and Protocols

#### Drosophila strains and husbandry

The fly stocks were maintained at a temperature of 25℃ under a 12-hour light and 12-hour dark cycle and were provided with a diet consisting of corn meal, yeast, and agar. For temperature-sensitive experiments, crosses were transferred to 18℃ incubators for eclosion. For the experiments with the daGS-Gal4 system, flies were raised on food with RU486 or vehicle after eclosion. For enoxacin treatment, parental fly crosses were settled on standard cornmeal added with drugs or vehicles until the progeny larvae conducted the crawling assay. *Drosophila* strains used in this study are listed in the Reagents and Tools Table. Unless otherwise specified, male flies were utilized for all experiments.

#### Locomotion tracing behavior and larval crawling assay

The adult fly and larval locomotor assay were carried out as described previously with minor modifications(Long *et al*., 2023). Adult males were quickly transferred to a 5 cm diameter open-field arena filled with silica gel (leaving a 5 mm space to the lid). Video recording captures adult flies’ free movement or third instar larvae for 1 minute with a high-resolution digital camera (see Reagents and Tools Table). Utilize the open-source Image J software combined with an animal tracker plugin to monitor and assess the movement patterns of fruit flies in the video. Convert the locomotion parameters from the video into digital format, including motion, immobility, and directional selection.

#### Eye imaging, retinal degeneration quantification, and scanning electron microscopy

Adult fly eyes were imaged with an Olympus SZX7 microscope with a SITUOLI STL- P830EISPM3C CMOS camera. Individual images were processed using Adobe Photoshop CS6 (Adobe). The compound eye pigmentation loss area was quantitatively analyzed using Image J software(Li *et al*, 2010). The surface area with disorganized ommatidia or necrotic part was considered as the degenerated area. The necrotic area was measured using Image J software, and the relative changes between groups were calculated.

For scanning electron microscopy (SEM), samples were gathered and fixed in a solution of 2.5% glutaraldehyde (#B46920, INNOCHEM, Inc.) in PBS and subsequently post-fixed for 20 min in 1.5% osmium tetroxide (#18465, Ted Pella Inc) in PBS. The samples were first dehydrated using ethanol with increasing concentrations of 30%, 50%, 70%, 80%, 90%, 95%, and 100% sequentially for 15 minutes each, followed by a 15-minute exposure to isoamyl acetate (10003128, Sinaopharm Group Chemical Reagent Co. LTD). Gold palladium coating was applied to the specimens attached to the metallic stubs and then analyzed with a HITACHI SU8100 scanning electron microscope.

#### RNA and miRNA sequencing

Total RNA was extracted from the cells or tissues using a Trizol reagent kit (Invitrogen, Carlsbad, CA, USA). RNA quality was assessed on an Agilent 2100 Bioanalyzer (Agilent Technologies, Palo Alto, CA, USA) and verified through RNase-free agarose gel electrophoresis. The total RNA was enriched using Oligo(dT) beads. Then, mRNA was fragmented into short fragments using a fragmentation buffer. The details were reverse transcribed into cDNA using NEBNext Ultra RNA Library Prep Kit for Illumina (NEB #7530, New England Biolabs, Ipswich, MA, USA). The double-stranded cDNA fragments were purified and end-repaired by adding a base. The ligation reaction was filtered with the AMPure XP Beads (1.0X). Ligated fragments were selected by size using agarose gel electrophoresis and polymerase chain reaction (PCR). The resulting cDNA library was sequenced using Illumina Novaseq6000 by Gene Denovo Biotechnology Co.

For miRNA sequencing, the RNA molecules in a size range of 18–30 nt were enriched by polyacrylamide gel electrophoresis (PAGE), after total RNA was extracted by Trizol reagent kit (Invitrogen, Carlsbad, CA, USA). The 3’ adapters were added, increasing the 36-48 nt RNAs. The 5’ adapters were then ligated to the RNAs as well. The ligation products were reverse transcribed, and the 140-160 bp size PCR products were enriched to generate a cDNA library and sequenced using Illumina HiSeq Xten by Gene Denovo (Biotechnology Co). mRNA, miRNA sequencing, and integrated analysis were performed using the OmicShare tools, an online platform (https://www.omicshare.com/tools).

#### Probe synthesis and electrophoretic mobility shift assay (EMSA)

Oligonucleotides of miRNA substrates were manufactured through in vitro transcription, utilizing T7 RNA polymerase (Cat #2017, Promega), with PCR-amplified templates and biotin-UTP (Perkin-Elmer). Biotin-labeled RNA transcripts (20 ng) were denatured for 5 minutes and incubated with 10 μg of protein extracts in 10 μl of Electrophoretic Mobility-Shift Assay Buffer (0.5 mg/ml bovine serum albumin, 40 mM KCl, 25 mM Tris-HCl pH 7.5, 1% TritongX-100, and 2 U/ml RNasin Plus RNase inhibitor) for 20 minutes at room temperature. After adding 4 ml of loading buffer (30% glycerol solution and 0.5% trypan blue), the sample was immediately run with a 6% acrylamide native gel. The gel was transferred to the NC membrane, stained with Striven (Invitrogen), and observed on a Gel DocTM EZ Imager system (Bio-Rad).

#### RNA and small RNA extraction and RT-qPCR

Small molecule RNA (miRNA, <200 nt) isolation from fly tissue or cultured cells (was conducted using a MiPure Cell/Tissue miRNA Kit (Vazyme). The cDNA template synthesis utilized the miRNA 1^st^ Strand cDNA Synthesis Kit (Vazyme) based on the stem-loop method, incorporating stem-loop primers (see Appendix Table S1) to elongate the template. Real-time PCR was carried out with a 7500 Fast Real-Time PCR System (Thermo Fisher Scientific). The fold induction of relative expression was determined using the ΔΔCT method, with snoR422 and GAPDH used as normalization controls in fly and cultured cell samples, respectively.

Total RNA was extracted using Trizol (Life Technologies) from cell culture, larvae, or adult, following the manufacturer’s instructions, including a DNase (Vezyme) treatment. cDNA was generated using the HiScript® III RT SuperMix kit (Vazyme) with 500 ng total RNA. Real-time PCR was carried out using ChamQ Universal SYBR qPCR Master mix (Vazyme) with the primers (see Appendix Table S2). Relative expression (fold induction) was calculated using the ΔΔCT method, and rp49 was used as a normalization control.

#### Cell culture and transfections and oxidative stress

HeLa or HEK293 cells were cultured in a standard Dulbecco’s modified Eagle’s/F-12 medium (DMEM), supplemented with 10% fetal bovine serum (Gibico) at 37℃ in an atmosphere of 5% CO2. The cell lines were authenticated by their respective sources and subsequently examined in our laboratory for Mycoplasma contamination.

Before transfection, the cells were examined for confluency and morphology, and the culture media was replaced. Cells were transfected with plasmid DNA expressing either TDP- 43 WT, A315T, or M337V with Lipofectamine 2000 (Thermo Fisher Scientific) transfection reagent according to the manufacturer’s procedure. Cells were maintained at 37℃ under a humidified atmosphere of 5% CO_2_ for 36 hours. For oxidative stress, cells were treated with 200 μM or 400 μM H_2_O_2_ for continuous culture for 4 hours before lysed for Western blot or with 4% PFA for immunostaining.

#### Western blot

Proteins lysate from cultured cell lines with RIPA buffer supplemented with complete mini protease inhibitor. The quantification of proteins was carried out using the BCA Protein Assay Kit (Thermo Fisher), and 300 μg of protein was then loaded onto pre-stained SDS-PAGE gels. The whole lysate of adult flies or larvae abdominal wall mixed with SDS sample loading buffer (200 mM Tris-HCl pH 6.8, 8% (w/v) SDS, 0.4% (w/v) BPB, 40% (v/v) glycerin, 4% (v/v) 2- ME) were heated at 95℃ for 5 min. Samples were added to the SDS-PAGE gel for electrophoresis with the electrophoresis system. Then, proteins were transferred to 0.45 μm PVDF membrane (Merck Millipore) using semi-dry or wet transfer. Unspecific binding was blocked using 5% non-fat skim milk or BAS in TBST for 15-30 min. Primary antibodies were diluted and incubated overnight at 4℃. According to the primary antibody, HRP-conjugated secondary antibodies (see key material table) were incubated at room temperature for 2 hours. Signals were developed using ECL Luminata^TM^ Western HRP Substrates (Merck Millipore) and the ChemiDoc system (Bio-Rad). To allow robust quantification, the exposure time for each blot was adjusted according to the expression level of an individual protein. The primary and secondary antibodies utilized in this study have been listed in the Reagents and Tools Table.

#### Acquisition and quantification of confocal images

For each experiment, the genotypes of interest were processed meanwhile. The images were obtained using the same parameters settings. For NMJ snaping, images of muscles 6 and 7 were acquired using the LSM Zeiss Software on a Zeiss LSM700 confocal microscope (63 × oil lens) and then analyzed using Fiji. Briefly, dissected adult brains and larvae were fixed in methanol for the primary antibody or 4% PFA for other antibodies. Three times washing of brain and NMJ using PBST (0.05% tween-20 in PBS) for 10 min. Then, the brain and larva were blocked in 10% goat serum (16210-064, Gibco) for 30 min. The primary antibodies were diluted and incubated with the tissue in the dark at 4℃ overnight. The second antibody was incubated at room temperature for 2 hours after three times washing. The primary and secondary antibodies utilized in this study have been listed in the Reagents and Tools Table.

#### Docking of enoxacin onto Dicer

Molecular docking of enoxacin of core ingredients and potential core targets of the Dicer protein was performed using AutoDock vina (see Reagents and Tools Table), and the results were visualized using Python molecule (PyMol) molecular graphics system.

#### Statistical analysis

All statistical analysis was performed with GraphPad Prism ver.9.0 (GraphPad, USA). Significance values for the dataset of a two-group comparison were calculated using a two-tailed unpaired *t*-test with Welch’s correction. For multiple comparisons, a one-way ANOVA followed by Tukey’s post-hoc test or two-way ANOVA followed by Bonferroni’s post-hoc test was conducted. All data have been presented as the Mean and the Standard Error of the Mean (SEM). Statistical significance was portrayed as *p < 0.05, **p < 0.01, ***p < 0.001, and ****p < 0.0001.

## Table of content

Appendix Table S1 2

Appendix Table S2 4

## Appendix Table S1

Oligonucleotides used for RT-qPCR to detect miRNA

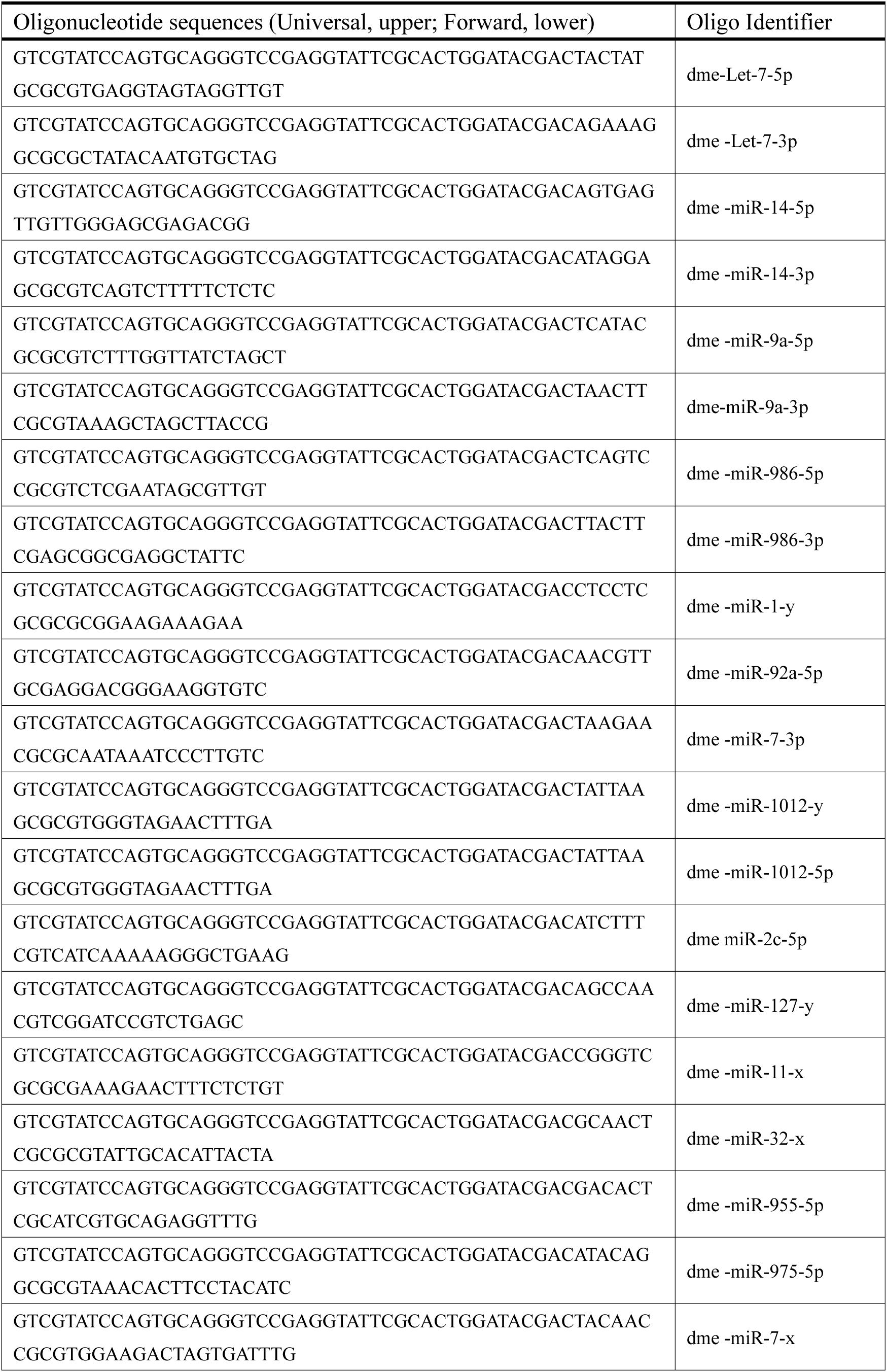
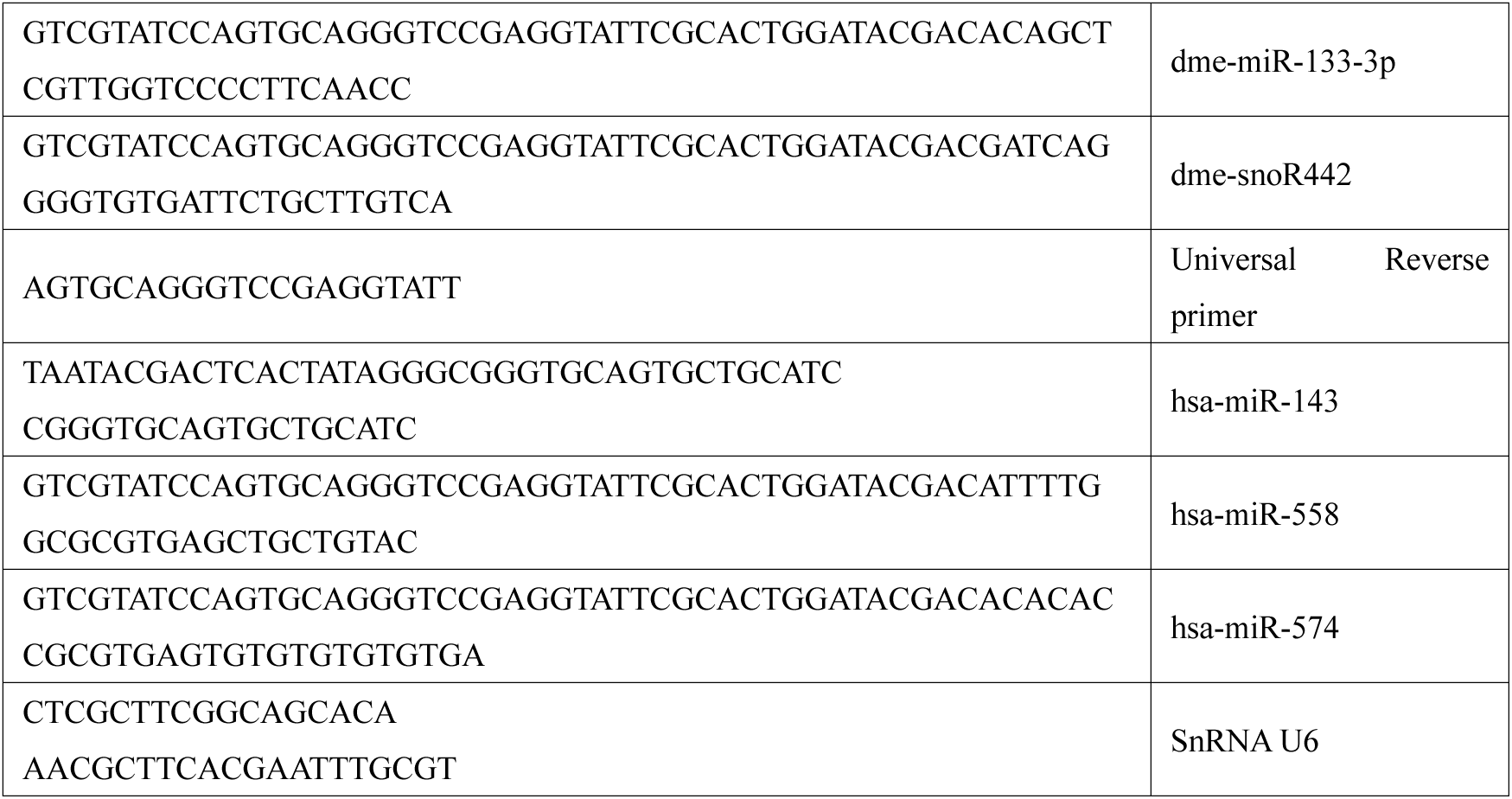

## Appendix Table S2

Primer design for RT-qPCR to detect mRNA in this study

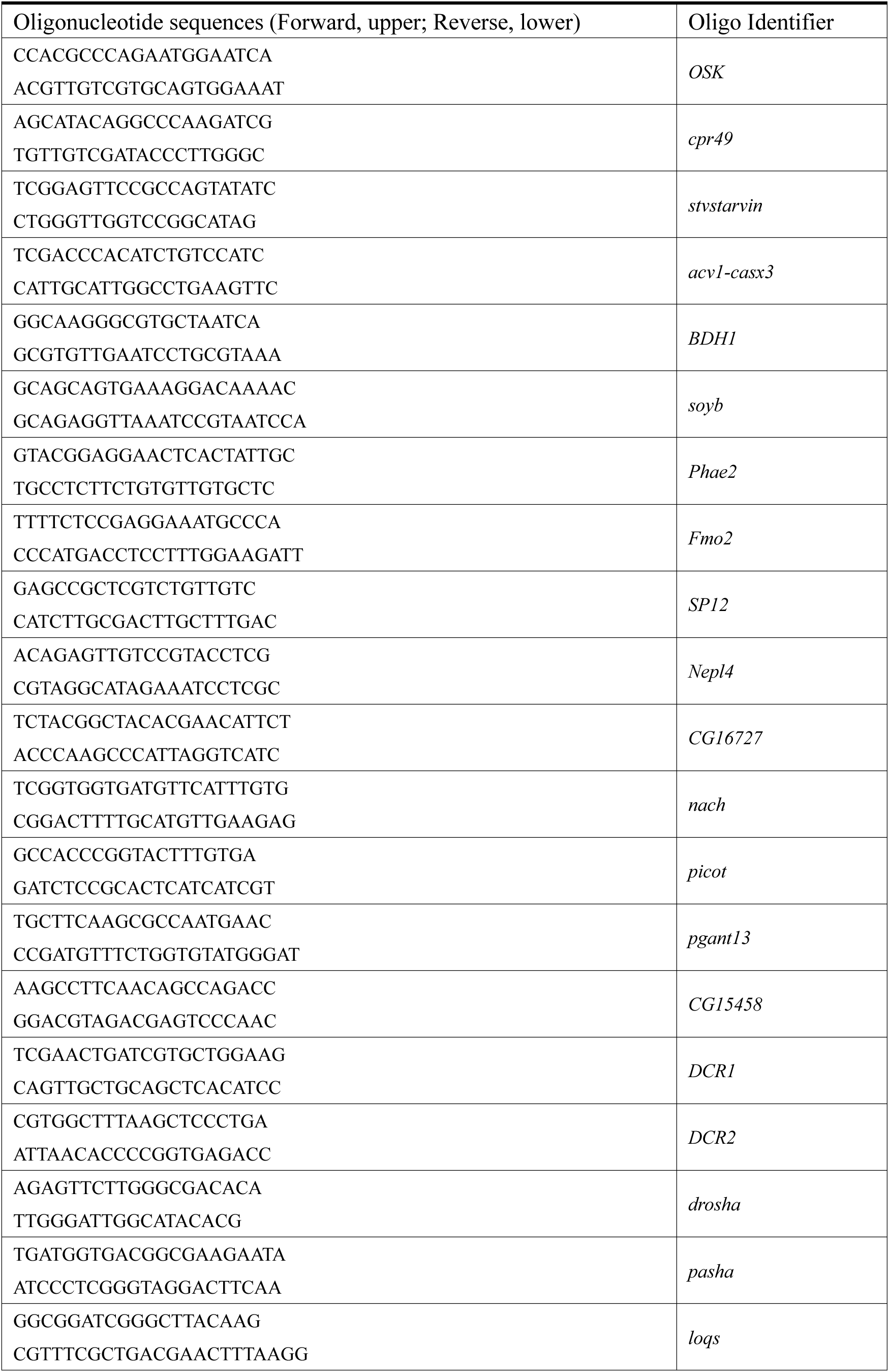
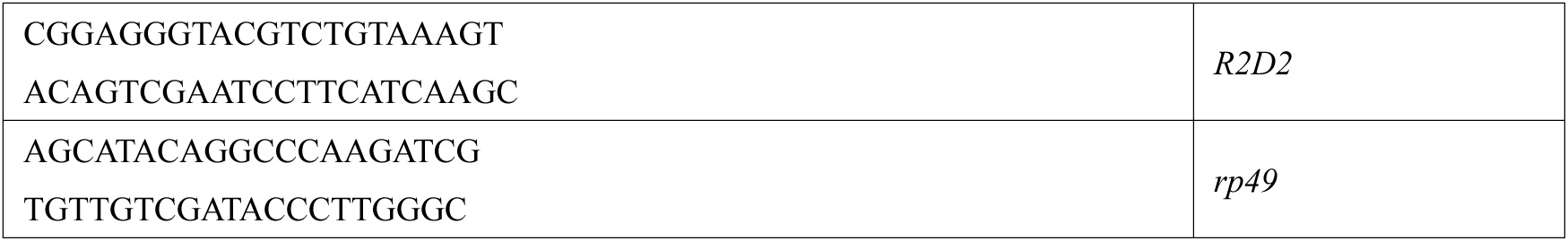

